# Constraint-based modelling revealed changes in metabolic flux modes associated with the Kok effect

**DOI:** 10.1101/2020.10.07.329854

**Authors:** Wei Qiang Ong, C. Y. Maurice Cheung

## Abstract

Constraint-based modelling was applied to provide a mechanistic understanding of the possible metabolic origins of the ‘Kok effect’ – the change in quantum yield of net photosynthesis at low light intensity. The well-known change in quantum yield near the light-compensation point (LCP) was predicted as an emergent behaviour from a purely stoichiometric model. From our modelling results, we discovered another subtle change in quantum yield at a light intensity lower than the LCP. Our model predicted a series of changes in metabolic flux modes in central carbon metabolism associated with the changes in quantum yields. We demonstrated that the Kok effect can be explained by changes in metabolic flux modes between catabolism and photorespiration. Changes in RuBisCO carboxylation to oxygenation ratio resulted in a change in quantum yield at light intensities above the LCP, but not below the LCP, indicating the role of photorespiration in producing the Kok effect. Cellular energy demand was predicted to have no impact on the quantum yield. Our model showed that the Kok method vastly overestimates day respiration – the CO_2_ released by non-photorespiratory processes in illuminated leaves. The theoretical maximum quantum yield at low light intensity was higher than typical measured values, suggesting that leaf metabolism at low light may not be regulated to optimise for energetic efficiency. Our model predictions gave insights into the set of energetically optimal changes in flux modes in low light as light intensity increases from darkness.

**One sentence summary:** The Kok effect can be explained by the changes in flux modes between catabolism and photorespiration.

## Introduction

The gross daytime CO_2_ production by a plant is traditionally thought to occur via the processes of photorespiration and day respiration. Day respiration is usually defined as the efflux rate of non-photorespiratory CO_2_ in illuminated leaves, expressed on a leaf area basis (Tcherkez et al., 2017b). Metabolic processes such as the tricarboxylic acid (TCA) cycle, the oxidative pentose phosphate pathway (OPPP) and all other non-photorespiratory decarboxylation reactions interact in a complex way that determines the rate of day respiration. Day respiration during photosynthesis had been studied for many years with multiple experimental techniques using classical gas exchange, isotopes or fluorescence (for reviews, see: (Tcherkez et al., 2012; Hodges et al., 2016; Tcherkez et al., 2017b); also see Tcherkez *et al.*, 2017b for a summary of techniques and its advantages and disadvantages). Due to its ease of implementation, the most popular technique used to study day respiration is the ‘Kok method’, which was developed by Bessel Kok and co-workers in 1948 where they discovered that the O_2_ evolution rate did not response linearly with light intensity in unicellular algae (Kok, 1948; Kok, 1949). The Kok method extrapolates the net CO_2_ assimilation rate to zero irradiance using data points above the break point to determine the day respiration, *R*_d_. Kok found that the O_2_ consuming respiratory flux in the dark was found to be higher than that at low light (Kok, 1948; Kok, 1949). This effect, now commonly referred to as the ‘Kok effect’, was interpreted as a consequence of dark respiratory inhibition by light. Further studies had demonstrated that the Kok effect is highly variable and is affected by environmental factors such as temperature (Dungan et al., 2003; Atkin et al., 2005; Ishii and Schmid Georg, 2014), concentration of CO_2_ and O_2_ (Ishii et al., 1977; Ishii et al., 1979; Sharp et al., 1984; Björkman and Demmig, 1987; Evans, 1987; Abadie et al., 2016; Farquhar and Busch, 2017), irradiance (Kok, 1948; Kok, 1949; Kok, 1956; Atkin et al., 1997), soil moisture (Illeris and Jonasson, 1999; Turnbull et al., 2001) and seasonality (Nordstroem et al., 2001; Wilson et al., 2001; Dungan et al., 2003; Kwon et al., 2009). A number of causes for the inhibitory effect of respiration by light had also been identified (see review: Heskel *et al.*, 2013).

While it is often easy to predict the rate of photorespiration using equations for gas exchange method and the internal CO_2_ mole fraction, estimating day respiration is more challenging as there is no equation that can predict its rate as a result of environmental parameters, CO_2_ mole fraction or photosynthesis. Despite being a difficult task, research over the past-half century has been trying to estimate *R*_d_ in the form of an equation or from a model of net carbon exchange (Hikosaka et al., 2016). However, fitting *R*_d_ into an equation might not be the best approach as there are many variables that affect it and the relationship between the variables are often complex and not well known.

Buckley & Adams presented a step forward in studying non-photorespiratory CO_2_ release at a systems level by developing a small flux balance model based on the balancing of coenzymes, ATP, NADH and NADPH, coupled with equations and variables from a photosynthesis model describing the processes generating and consuming the coenzymes (Buckley and Adams, 2011). The Kok effect was built into the Buckley & Adams model based on the assumption that the OPPP was inhibited indirectly by light. In this study, we used a large-scale stoichiometric metabolic model to explore the multiple metabolic processes that could contribute to the Kok effect. The main advantage of our model is that it only contains stoichiometric information of reactions in central metabolism, and the Kok effect was predicted as an emergent property of the model. In addition, since our model contains individual reactions covering in the central metabolism of a C_3_ leaf, we were able to capture a complete picture of the metabolic processes that resulted in the Kok effect, which are typically difficult to study experimentally with small fluxes at low light intensity.

## Results

### The Kok effect was predicted based on a purely stoichiometric model

Using a charge- and mass-balance stoichiometry model of core plant metabolism extracted from Shameer et al., (2018), the light response curve was predicted based on the constraints of fixed phloem output and maintenance costs and the primary objective function of minimising photon influx (see Material and Methods for more details) with the ratio of ribulose bisphosphate carboxylase/oxygenase (RuBisCO) carboxylation to oxygenation reactions set to 3:1 as in previous studies (Cheung et al., 2014; Cheung et al., 2015; Shameer et al., 2018). The predicted light response curve of net CO_2_ flux of a C_3_ leaf under different incident photon intensity showed a break at a low photon intensity of about 20 μmol m^−2^ s^−1^ (Fig. 1). Coincidently, at this break point, the net CO_2_ flux is zero and this point is also commonly referred to as the light compensation point (LCP) where the CO_2_ consumed during photosynthesis equals the CO_2_ released during respiration. LCP is also commonly used as the reference point to distinguish the dark and the low light period. The region of no light to LCP (i.e. in this case, 0 – 20 μmol m^−2^ s^−1^) is classified as the dark / night period and is assigned as stage 1 and the region thereafter is classified as the low light / day period and is assigned as stage 2 (Fig. 1). The intercepts of the resulting linear fits for the points in stage 1 and stage 2 represent the apparent rates of respiration in the dark (night respiration, *R*_n_) and in the day (day respiration, *R*_d_) respectively. *R*_n_ and *R*_d_ were predicted to be 2.98 μmol m^−2^ s^−1^ and 1.44 μmol m^−2^ s^−1^ respectively. It is worth noting that the break in linearity of the light curve was predicted from a purely stoichiometric model, suggesting that the Kok effect can be explained by stoichiometry alone without any processes involving kinetics. Besides the break in linearity of the net CO_2_ flux curve at LCP, another subtle change in the gradient of the curve was observed to have occurred in stage 1, at about 12 μmol m^−2^ s^−1^ (Fig. 1b). A similar break can be observed in the light response curve for oxygen release (Fig. S1). This break was assigned as Point A and its significance was highlighted in the subsequent sections with changes in flux modes around Point A.

**Fig. 1.**
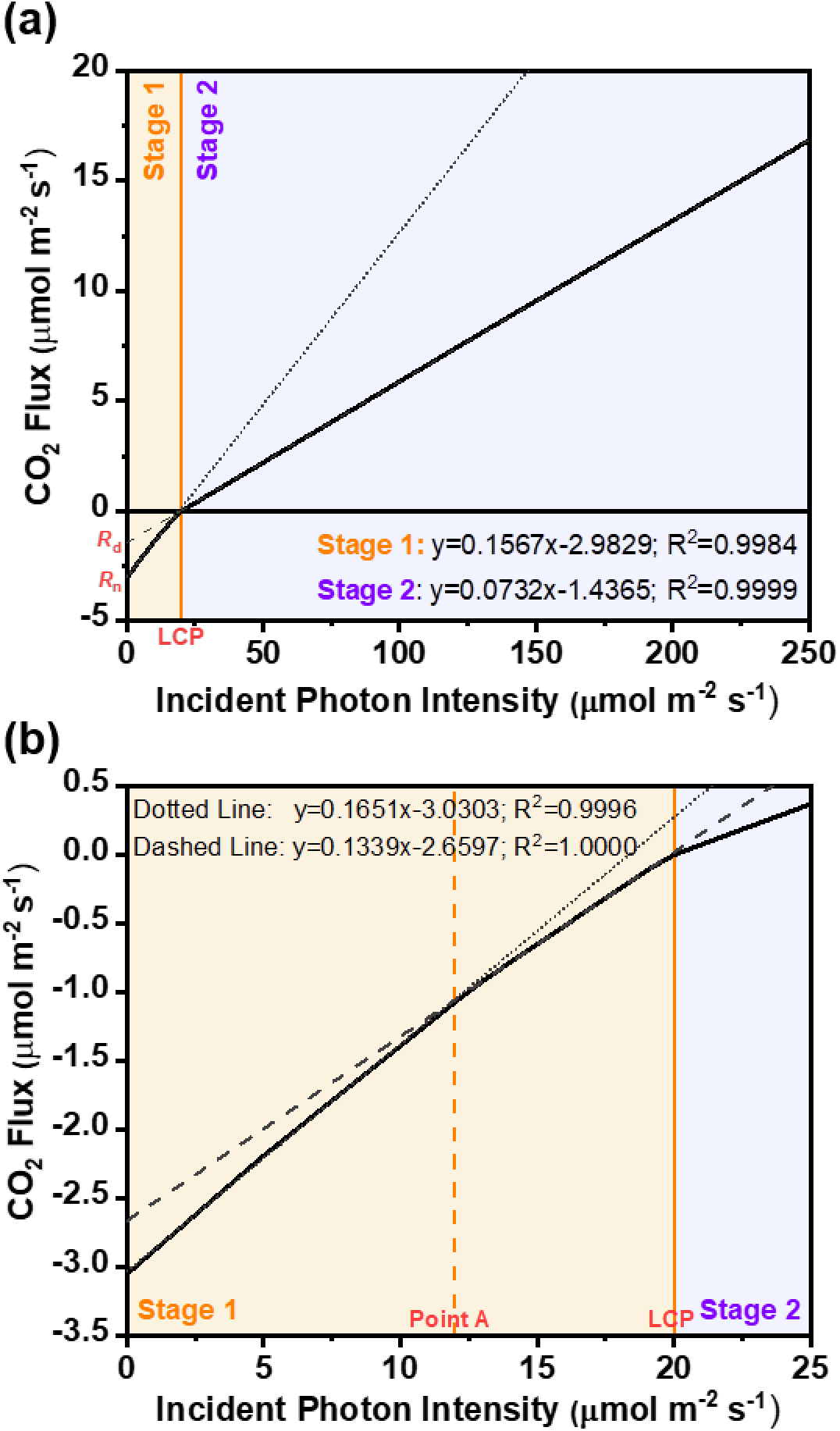
Predicted light response curves of CO_2_ flux of C_3_ leaf under different incident photon intensity. The ratio of carboxylation to oxygenation reactions by RuBisCO was fixed at 3:1. In Fig. 1a, at low incident photon intensity, a break in the linear light response curve was observed to occur at an incident photon intensity of about 20 μmol m^−2^ s^−1^, which coincides with the light compensation point (LCP). The resulting linear fits for the points below the LCP (stage 1) and above the LCP (stage 2) were plotted as a dotted and dashed line respectively. The estimations of respiration rate of the C_3_ leaf in the day (*R*_d_) and in the dark (*R*_n_) by the Kok method were given by the y-intercept of the dashed line and dotted line respectively and from the graph. *R*_d_ and *R*_n_ were found to be 1.44 μmol m^−2^ s^−1^ and 2.98 μmol m^−2^ s^−1^ respectively. Fig. 1b is an expanded region of the light response curve from 0 – 25 μmol m^−2^ s^−1^. Another change in the gradient of the light response curve at about 12 μmol m^−2^ s^−1^ (Point A) was observed. The linear fit line for the data from 0 μmol m^−2^ s^−1^ to Point A was represented by a dotted line with an equation of y = 0.17x – 3.03. The linear fit line for the data from 13 to 19 μmol m^−2^ s^−1^ was represented by a dash line with an equation of y = 0.13x – 2.66.

### Photorespiration affects *R*_d_ but not *R*_n_ from the light curve

In order to understand the effect of photorespiration on the Kok effect, we repeated our simulations with different RuBisCO carboxylation to oxygenation ratios ranging from 0.5:1 to 100:1. Our results showed that decreasing the ratio led to a more moderate slope (lower gradient) in stage 2, i.e. lower quantum yield (amount of CO_2_ fixed per photon) (Fig. 2). This is expected since a decrease in the RuBisCO carboxylation to oxygenation ratio means an increase in photorespiration and thus more light will be needed to fix the same amount of CO_2_ as photorespiration is an energy-consuming process. Under atmospheric gaseous conditions, RuBisCO has a carboxylase/oxygenase activity ratio range of 1.86:1 to 3:1 (Abadie et al., 2016). Based on this range of RuBisCO carboxylase/oxygenase ratio, *R*_d_ obtained from our light response curve (Fig. 2) was estimated to be in the range of 1.11–1.44 μmol m^−2^ s^−1^, which was lower than *R*_n_ by 52–63%. We noted that these estimates were higher than the well accepted inhibition of leaf respiratory metabolism by light of 20–40% (Tcherkez et al., 2017a). In particular, our model overestimated the quantum yield of stage 1. The theoretical quantum yield of stage 1 (~0.16; Fig. 1) is higher than the typically observed quantum yield under standard conditions (~0.1; (Tcherkez et al., 2017a)), suggesting that the regulation of leaf metabolism in stage 1 may not be optimised for energy efficiency. It should be noted that at high RuBisCO carboxylase to oxygenase ratio, the quantum yield of stage 2 can get so high that the Kok effect (changes in quantum yields between stage 1 and stage 2) was close to disappearing (Fig. 2), suggesting that low oxygen and/or high CO_2_ conditions could lead to the disappearance of the Kok effect. We observed that when the ratio of RuBisCO’s carboxylation/oxygenase activity was 2.5:1, the predicted quantum yield (i.e. gradient of linear fitted line in stage 2 = 0.067) was very similar to the experimental result reported by Farquhar and Busch (2017), indicating a RuBisCO carboxylase to oxygenase ratio of 2.5:1 would be a more appropriate constraint to model the physiological condition. At this ratio, the model predicted *R*_d_ and *R*_n_ to be 1.32 and 2.98 μmol m^−2^ s^−1^ respectively using the Kok method. All subsequent simulations and analyses were performed using a RuBisCO carboxylase to oxygenase ratio of 2.5:1 to investigate the various metabolic processes in C_3_ leaf that give raise to the Kok effect. We also noted that changing the ratio of RuBisCO carboxylation to oxygenation activities did not have any effect on the gradient and y-intercept of the light response curve in stage 1, which means *R*_n_ is not affect by photorespiration. Inspection of the predicted metabolic fluxes showed that RuBisCO and glycine decarboxylase complex (GDC), the enzyme complex involved in photorespiration in the mitochondrion, only became active at photon intensities above the LCP in all simulated scenarios (Fig. 3), which explains why photorespiratory rate has no effect on *R*_n_. The increase in RuBisCO flux at the LCP indicated that Calvin-Benson cycle was predicted to only activate at light intensities higher than the LCP. Given that the objective function used optimises for maximum energy use efficiency, our results suggest that it is energetically optimal for RuBisCO to only be active as light intensity is higher than the LCP.

**Fig. 2.**
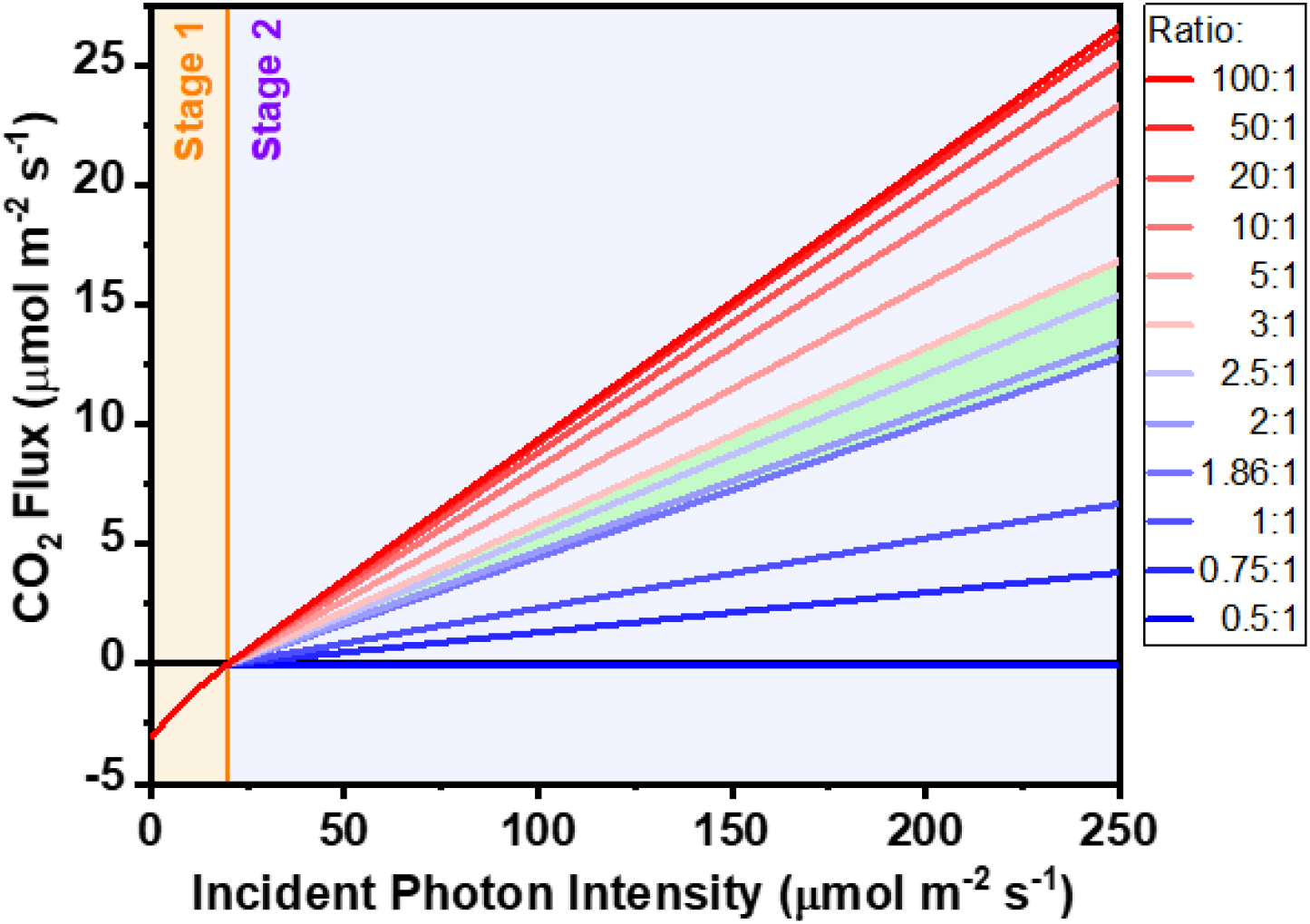
Predicted light response curve at different ratios of RuBisCO carboxylase to oxygenase activity. The area highlighted in green showed the typical range of RuBisCO carboxylation to oxygenation ratio under atmospheric gaseous conditions. When the incident photon intensity was below LCP (20 μmol m^−2^ s^−1^; orange vertical line), the CO_2_ fluxes were the same for all the different ratios. Only when the incident photon intensity increased above LCP, an increase in the ratio of RuBisCO carboxylase to oxygenase activity led to an increase in CO_2_ production rate and the quantum yield of stage 2.

**Fig. 3.**
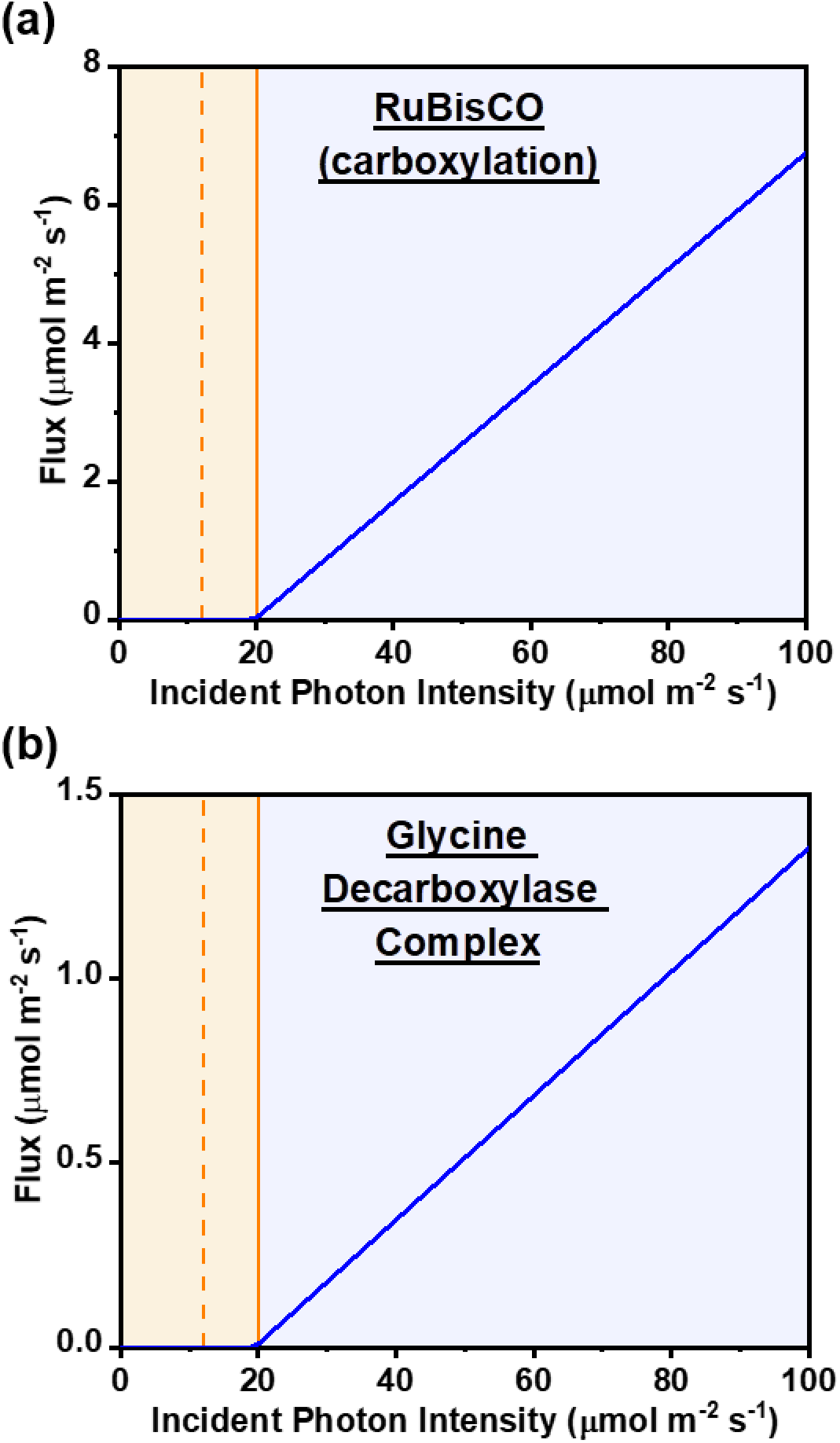
Fluxes of RuBisCO carboxylase (Fig. 3a) and glycine decarboxylase complex (GDC) (Fig. 3b) under different incident photon intensity. Ratio of RuBisCO carboxylase/oxygenase activity ratio was fixed 2.5:1. The graphs showed that RuBisCO carboxylase, involved in photosynthetic carbon fixation, and GDC, involved in the photorespiratory pathway, were predicted to be active only at light intensities above the LCP. Point A and LCP are represented by an orange vertical dashed and solid lines respectively.

### Down-regulation of respiratory decarboxylation as light intensity increases from darkness

To understand the metabolic mechanism that gives raise to the Kok effect, we inspected the representative reactions involved in the generation of CO_2_ (Fig. 4 & Fig. S2). In general, most of these reactions can be classified into two groups. The first group of reactions had a constant metabolic flux as light intensity increased and their absolute flux value were generally very small (Fig. S2). These reactions are generally involved in biomass synthesis. On the other hand, the second group of reactions showed a change in the rate of activity at either around Point A (e.g. formate dehydrogenase, oxoglutarate dehydrogenase and mitochondrial pyruvate dehydrogenase) or at the LCP (e.g. cytosolic glucose 6-phosphate dehydrogenase) as light intensity increased. The respiratory decarboxylation enzymes in the second group were involved in either the OPPP, TCA cycle, Calvin cycle, photorespiratory pathway or ammonium assimilation, and the cumulative fluxes of these enzymes reactions showed a decreasing trend in stage 1 (Fig. 4), indicating that (1) these metabolic processes could play a role in contributing to the Kok effect based on their predicted changes in fluxes as light intensities vary and (2) these enzymes are predicted to be downregulated as light intensity increases from dark to low light.

**Fig. 4.**
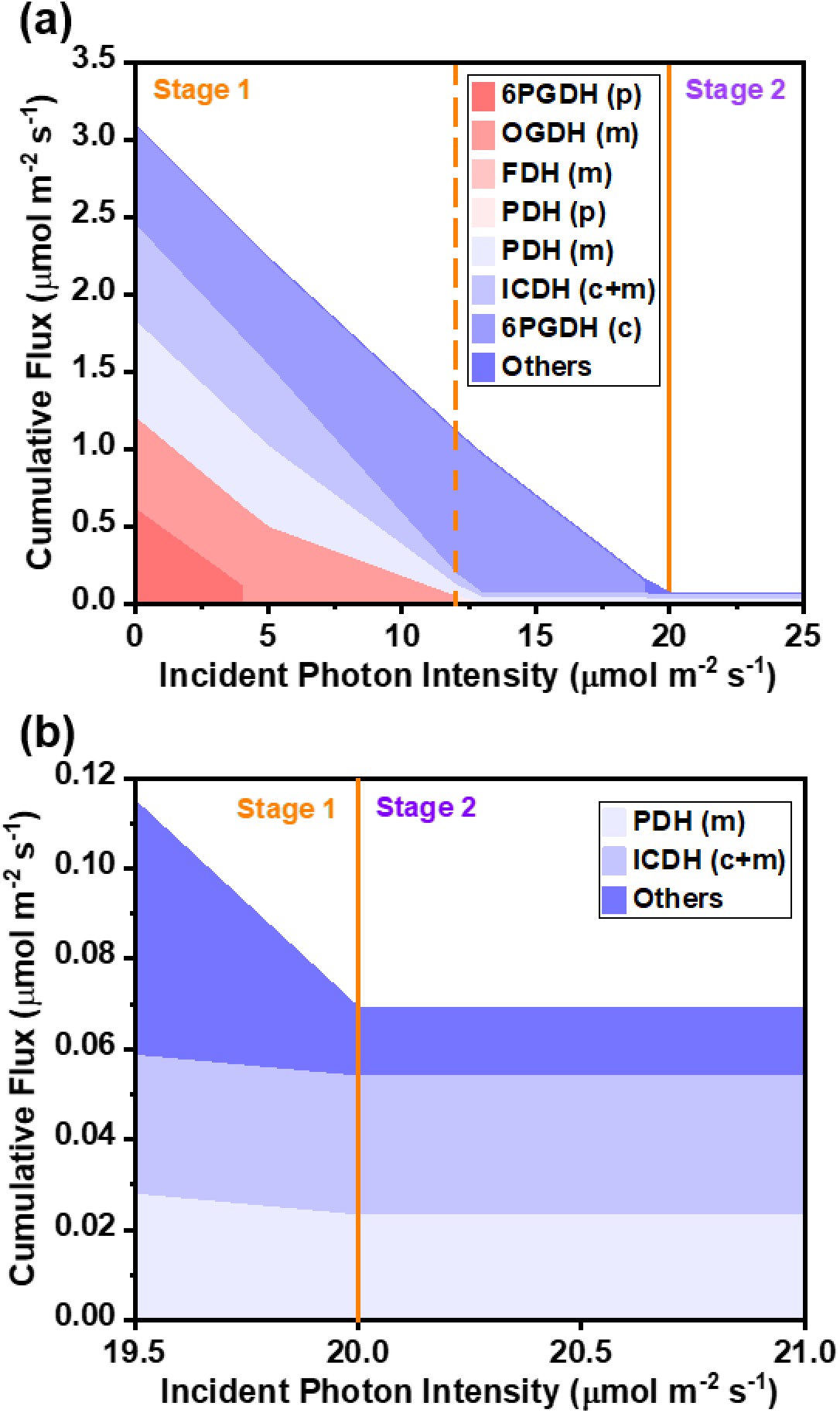
Cumulative fluxes of enzymes involved in carbon dioxide producing reactions under different incident photon intensity. The cumulative graph in Fig. 4a showed that the fluxes of these CO_2_-generating enzymes decrease with increasing photon intensity and plateau off at light intensities above the LCP. The decreasing trend can mostly be attributed to 6PGDH (c), OGDH, PDH (m) and ICDH as their fluxes showed large changes with changes in photon intensity. Other CO_2_-producing enzymes gave very small constant fluxes (refer to Fig. S2 in Supplemental Data for the individual flux of these enzymes). Fig. 4b showed an expanded region of the light curve from 19.5 – 21.0 μmol m^−2^ s^−1^. As shown in the graph, after LCP, only PDH (m), ICDH and ‘Others’ were active, but their fluxes were very small compared to the CO_2_ release in darkness. Their cumulative fluxes was only 0.07 μmol m^−2^ s^−1^ (~2.3% of the initial cumulative fluxes at zero photon intensity). Abbreviation: 6PGDH: 6-Phosphogluconate dehydrogenase; OGDH: 2-oxoglutarate dehydrogenase; FDH: Formate dehydrogenase; PDH: Pyruvate dehydrogenase; ICDH: Isocitrate dehydrogenase. Acetolactate synthase, Arogenate dehydrogenase, Carboxycyclohexadienyl dehydratase, Diaminopimelate decarboxylase, Formate dehydrogenase, Glutamate decarboxylase, Imidazole glycerol phosphate synthase, Prephenate dehydrogenase had very small constant fluxes and were group together in ‘Others’. Lowercase letters c, m and p in parenthesis stand for isozymes in the cytosol, mitochondrion and plastid respectively. LCP and Point A are represented by an orange vertical solid and dashed line respectively.

The OPPP generates NADPH in successive oxidative reactions when converting glucose 6-phosphate into ribulose 5-phosphate. There had been reports that suggested that the one of the factors that led to the Kok effect was the suppression of the OPPP in low light (Singh et al., 1993; Farr et al., 1994; Huppe et al., 1994; Buckley and Adams, 2011), which is consistent with our modelling results. Our simulations showed that the OPPP were only active in stage 1 and not in stage 2 (Fig. 5). As the intensity of the incident photons increased, the OPPP flux decreased, along with the amount of CO_2_ generated, suggesting that the role of OPPP was primarily to produce reducing equivalents, in the form of NADPH, in the dark (< 20 μmol m^−2^ s^−1^) before it is taken over by the photosynthetic light reactions.

**Fig. 5.**
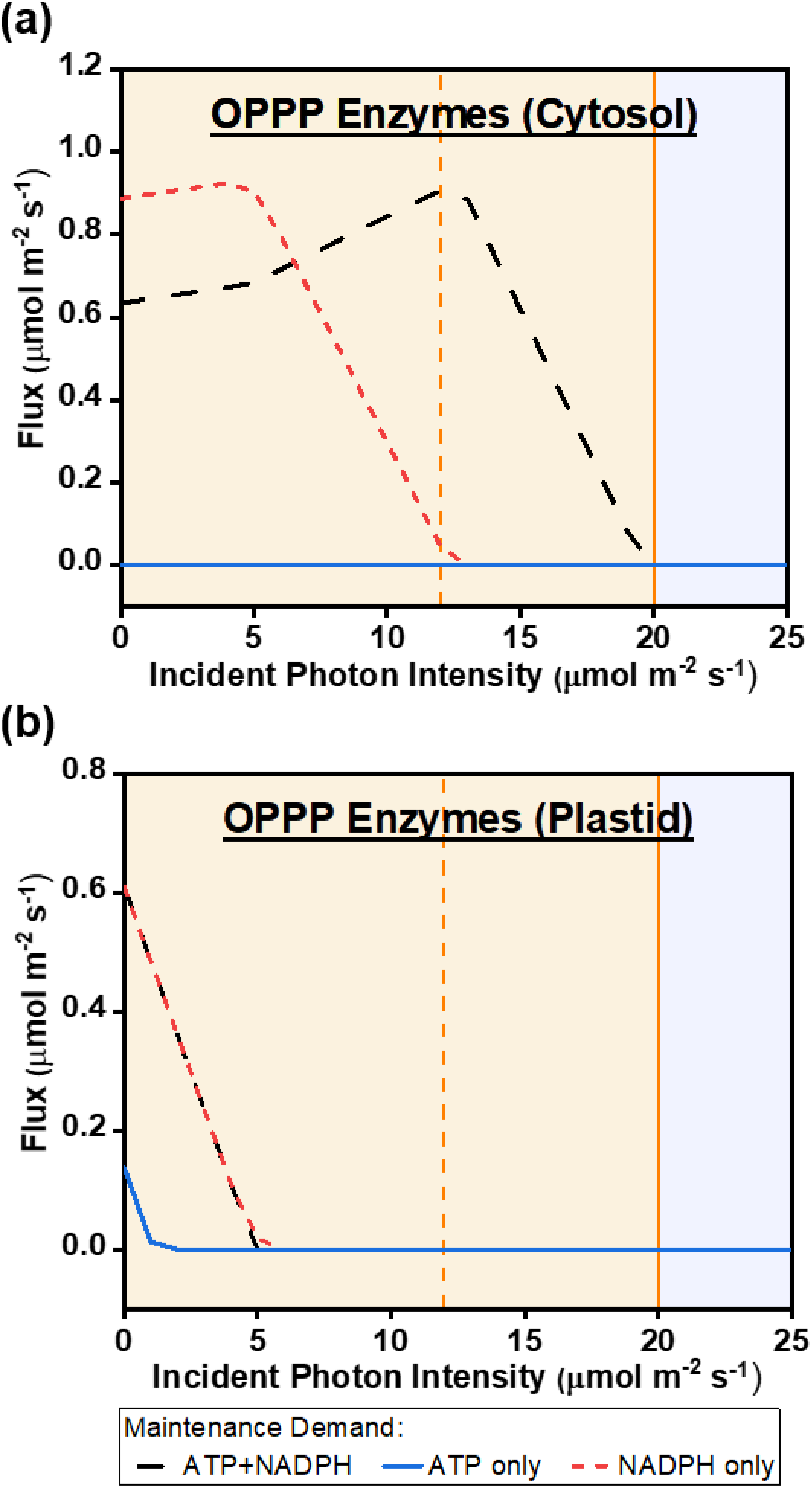
Predicted fluxes of the cytosolic (Fig. 5a) and plastidial (Fig. 5b) oxidative pentose phosphate pathway (OPPP) at different incident photon intensities. Identical fluxes were obtained for all three enzymes, glucose-6-phosphate dehydrogenase, gluconolactonase and 6-phosphogluconate dehydrogenase, in the same modelled conditions. The black curve showed the flux obtained under normal modelling constraints where the maintenance demand of both ATP and NADPH were present. The blue curve represented the flux where there was only ATP demand and no NADPH demand (i.e. NADPH demand was fixed at 0 μmol m^−2^ s^−1^). Vice versa, the red curve represented the flux where the ATP demand was fixed at 0 μmol m^−2^ s^−1^ while maintaining the NADPH demand. Point A and LCP are represented by an orange vertical dashed and solid lines respectively.

There had been previous experimental studies reporting the decrease in TCA enzymes abundance or activities in the mitochondria during the transition from dark to light (Budde and Randall, 1990; Tovar-Méndez et al., 2003; Lee et al., 2010). Our model predictions showed the same observations. Except for malate dehydrogenase, which is involved in redox shuttles, the predicted fluxes of all other TCA cycle reactions and pyruvate dehydrogenase decreased to zero or close to zero with light intensity above Point A (Fig. S3). This suggested that one of the factors that resulted in a metabolic transition at Point A was the TCA cycle. As pyruvate dehydrogenase, isocitrate dehydrogenase and oxoglutarate dehydrogenase are involved the generation of CO_2_, a decrease in fluxes through these reactions from dark to low light led to a decrease in CO_2_ generation. Note that the isozymes of isocitrate dehydrogenase are often involved in shuttling reducing power between the cytosol and the mitochondrion. In our model prediction, only the NADP^+^-dependent isocitrate dehydrogenase (ICDH) was active, and the net flux for the cytosolic and mitochondrial ICDH were reported throughout this study. Based on the results from our purely stoichiometric model, it can be concluded that it is energetically optimal to downregulate decarboxylating enzymes in the OPPP and the TCA cycle as light intensity increases from zero to the LCP.

### The effects of cellular maintenance on the Kok effect

One of the main functions of the TCA cycle and the OPPP, the processes predicted to play a role in the Kok effect, is to produce energy and reducing power, in the form of ATP and NADPH, to support cellular maintenance processes. And hence, we repeated the analysis with a range of ATP and NADPH maintenance costs to determine how the energetics of cellular maintenance affects the Kok effect. From our simulation results, we observed a shift in the light response curve to the right as we increased ATP and NADPH maintenance costs (Fig. 6). When the maintenance costs were increased by 5– folds, the LCP changed from 20 μmol m^−2^ s^−1^ to 90 μmol m^−2^ s^−1^ and *R*_d_ increased from 1.44 μmol m^−2^ s^−1^ to 6.09 μmol m^−2^ s^−1^. Interestingly, the ratio of *R*_d_ to *R*_n_ remained close to constant (0.44–0.45), i.e. *R*_d_ is lower than *R*_n_ by 55–56%, regardless of the magnitude of the maintenance costs. In summary, our results predicted an increase in *R*_d_, *R*_n_ and the LCP as maintenance energy demands increases, while the quantum yields of phase 1 and phase 2 remained close to constant.

**Fig. 6.**
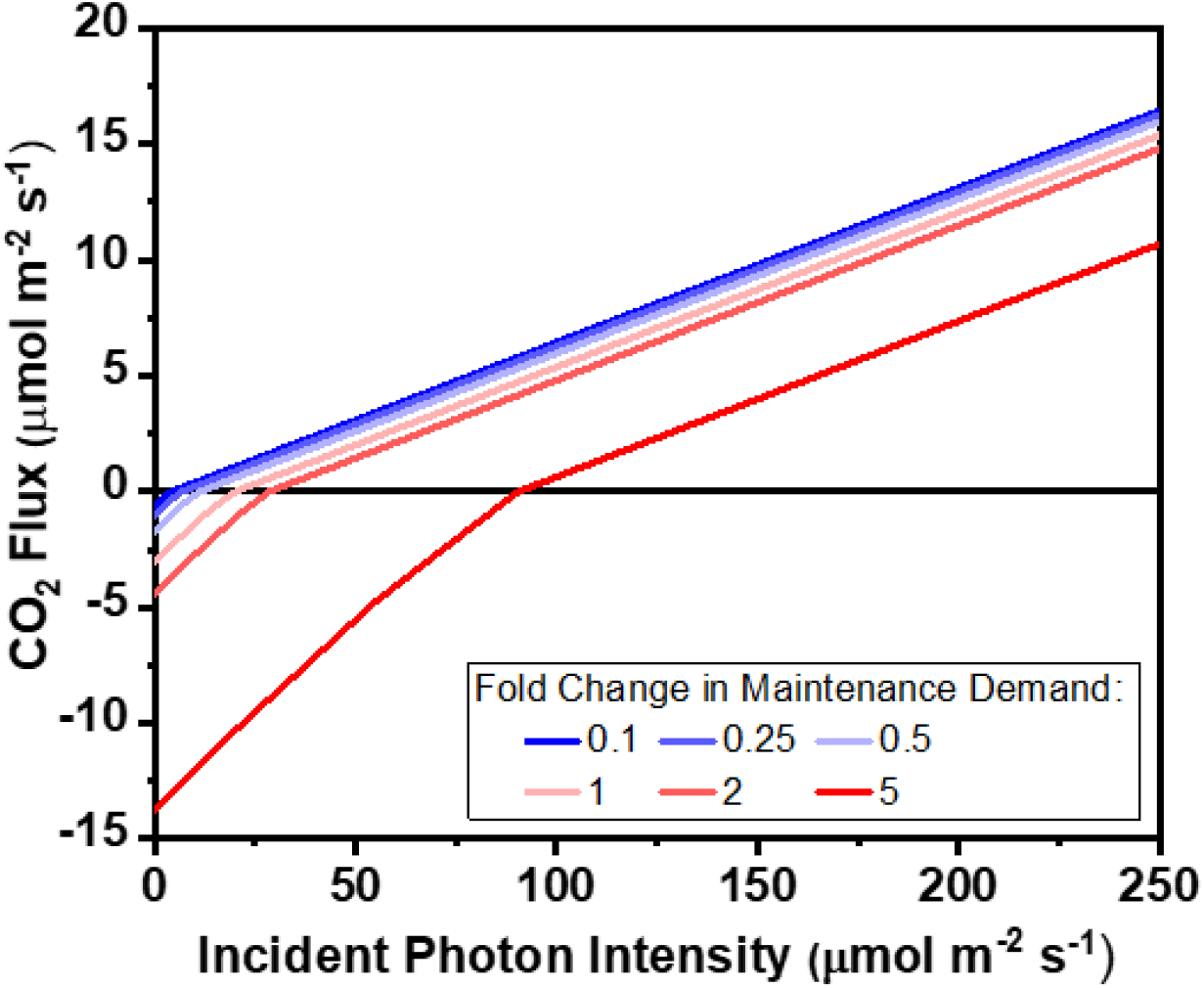
Predicted light response curve under different fold changes of ATP and NADPH maintenance demands. Increasing fold changes in the magnitude of maintenance demands were shown as different colours from blue to red. Changing the maintenance costs did not have any effect on the quantum yields at low light as the gradients of the linear fit lines remained the same for all maintenance demands modelled.

Subsequently, we investigated the individual effects of ATP and/or NADPH had on the Kok effect independently. As shown in Fig. 7, when the demand for either ATP, NADPH or both was fixed at zero, the photosynthetic light curve shifted to the left as expected due to a decrease in energy demand. When there was only a demand for NADPH but not ATP for maintenance processes in the leaf cell, the LCP was about 13 μmol m^−2^ s^−1^. However, when NADPH demand was zero and there was only a demand for ATP for maintenance, the LCP dropped by 50% from 20 μmol m^−2^ s^−1^ to 10 μmol m^−2^ s^−1^. The LCP further dropped to about 2.5 μmol m^−2^ s^−1^ when there was no maintenance demand for ATP or NADPH in the cell. These results indicated that the CO_2_ released in stage 1 were mostly driven by the production of ATP or NADPH needed for maintenance processes. This was consistent with what we had observed earlier where the enzymatic fluxes for metabolic processes such as the OPPP and the TCA cycle were only upregulated in stage 1. We next looked at the individual effects of ATP and NADPH maintenance on the individual enzymatic fluxes of the metabolic processes that were related to day respiration to investigate how these processes affected the Kok effect.

**Fig. 7.**
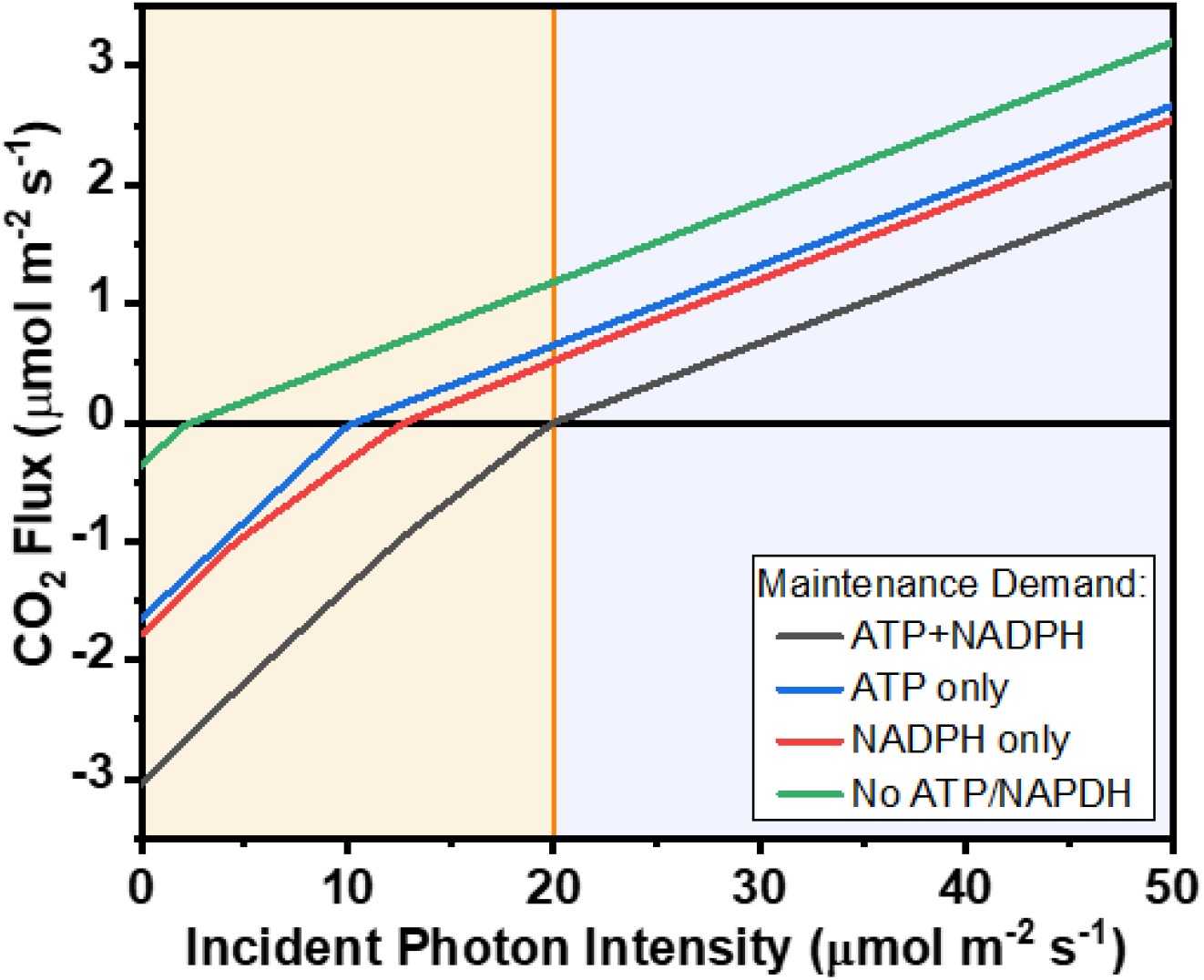
Predicted light response curves in scenarios with different types of energetics demand for cellular maintenance. Scenarios with both ATP and NADPH maintenance demands, ATP demand only, NADPH demand only and no maintenance demands were shown in black, blue, red and green lines respectively. LCP is represented by an orange vertical solid lines.

#### (A) Effects of ATP maintenance on metabolism in low light

The lower glycolysis pathway consists of reactions involving 3-carbon molecules, where glyceraldehyde 3-phosphate (GAP) is being converted into pyruvate. The production of ATP is being carried out by phosphoglycerate kinase and pyruvate kinase in this pathway. In effect, it had been shown that the activity of pyruvate kinase is higher in the dark than in the light and the production of pyruvate is inhibited by light (Lin et al., 1989; Scheible et al., 2000). In the standard scenario with both ATP and NADPH maintenance, our results showed that the enzymatic fluxes of lower glycolysis pathway were upregulated in stage 1 and the fluxes dropped in a steady fashion to Point A as light intensity increased (Fig. S4), suggesting that the lower glycolysis pathway, which is responsible for the production of ATP, were only upregulated before Point A and not the whole Stage 1. And as expected, when there was only a demand for ATP maintenance only but not NADPH, the lower glycolytic flux dropped only marginally as compared to the standard scenario. In contrast, a more than 80% drop in fluxes was observed when there was a demand for NADPH only.

The TCA cycle is also involved in the production of ATP by producing NADH for the mitochondrial electron transport chain. The results of the effects of ATP and NADPH on the fluxes of TCA enzymes were shown in Fig. S3. In general, for most TCA enzymes, a notably decrease in fluxes was observed when there was only a demand for NADPH and not ATP. The initial drop in absolute value was about 80% and the curves flatten out at Point A, suggesting that the role of the TCA cycle was primarily to produce the ATP needed in maintenance till Point A. After Point A, the production of ATP in the leaf will be taken over by other metabolic processes, mainly the photosynthetic light reaction. When there was only a demand for ATP, the fluxes through most reactions in the TCA cycle, except malate dehydrogenase which is usually involved in redox shuttles, dropped marginally compared to the scenario with both ATP and NADPH maintenance demands (Fig. S3), as expected given the role of the TCA cycle in ATP production.

#### (B) Effects of NADPH maintenance on metabolism in low light

The primarily role of the OPPP to provide NADPH for cellular maintenance is well known. As shown in Fig. 5, the OPPP enzymes were only active in Stage 1 and when there is a demand for NADPH. When there was only an ATP demand in the leaf, no cytosolic fluxes was observed and the OPPP flux in the plastid dropped significantly, by about 70%, as compared to the normal condition when there was a demand for both ATP and NADPH. And the plastid OPPP flux disappeared almost immediately upon increasing light intensity from completely darkness, at an extremely low light intensity of 2 μmol m^−2^ s^−1^. On the other hand, when there was only a demand for NADPH in the leaf, the initial enzymatic flux was higher (0.89 μmol m^−2^ s^−1^) than with both ATP and NADPH maintenance (0.63 μmol m^−2^ s^−1^). However, as light intensity increased, the OPPP flux decreased reached zero at around Point A, instead of at the LCP for the scenario with ATP and NADPH maintenance. It is also interesting to note that with both ATP and NADPH maintenance, the OPPP flux had a local peak at around Point A, which was absent for the scenarios with only ATP or only NADPH. This suggests that there could be interactions between the production of ATP and NADPH in stage 1, possibly involving lower glycolysis.

A variant of the lower glycolysis pathway involves the direct conversion of GAP to 3-phosphoglycerate by glyceraldehyde-3-phosphate dehydrogenase (NADP^+^) (GAPN). GAPN uses NADP^+^ as the electron acceptor and does not produce ATP. Unlike the other enzymes in the lower glycolysis pathway, GAPN was predicted to only be active with light intensity higher than Point A in the standard scenario, and as the light intensity increased, the flux increased before reaching a maximum and plateauing off at the LCP (Fig. 8). Similarly, unlike the other lower glycolysis enzymes, when there was no NADPH demand in the leaf, GAPN remained totally inactive and the flux remained at 0 μmol m^−2^ s^−1^ as light intensity increased, suggesting that the function of this enzyme was solely to produce NADPH needed for maintenance. When there was only a demand for NADPH in the leaf, the enzyme became active much earlier and reached a maximum at Point A (Fig. S4f).

**Fig. 8.**
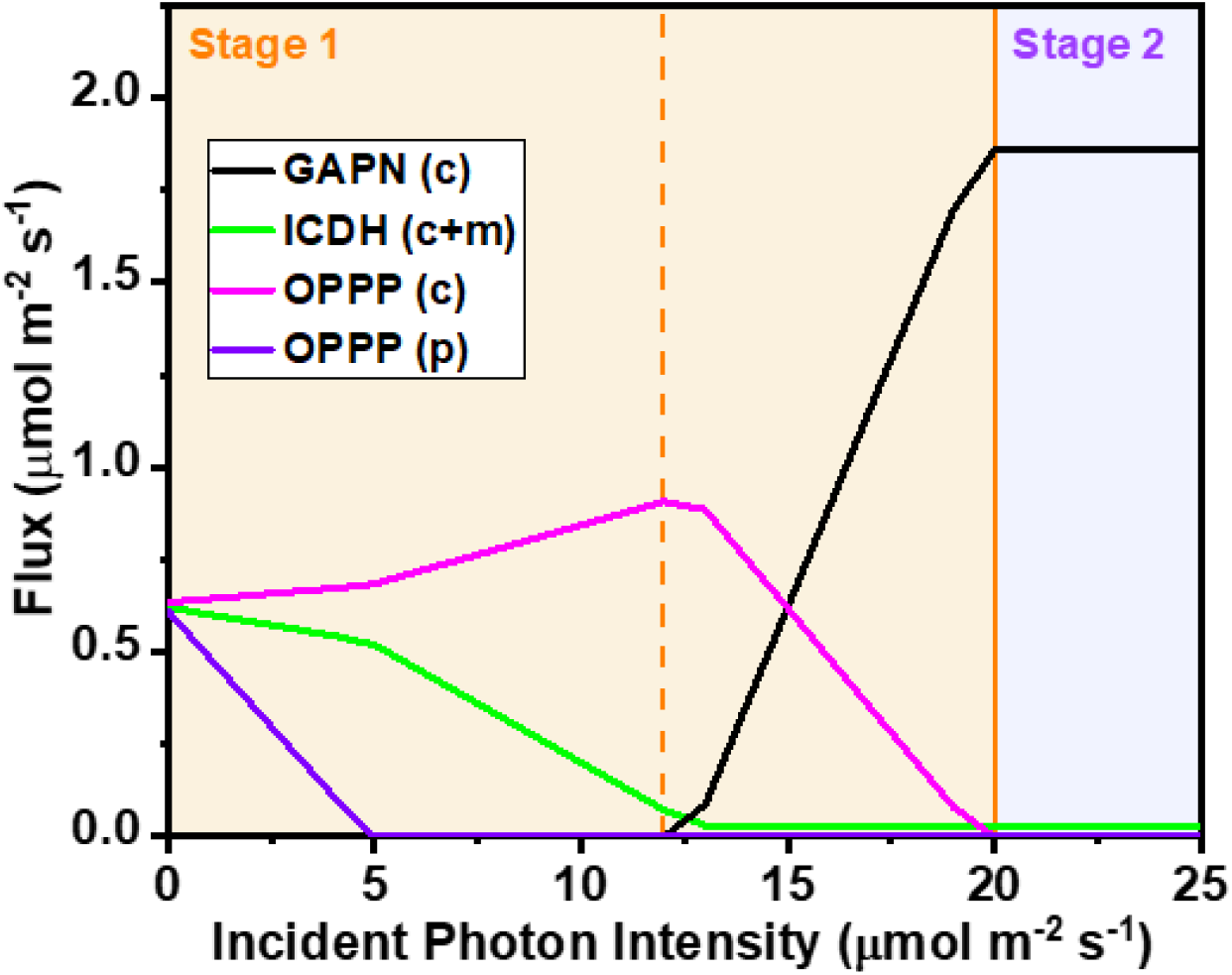
Fluxes of NADPH-generating enzymes, glyceraldehyde 3-phosphate dehydrogenase (NADP^+^) (GAPN) (black), isocitrate dehydrogenase (ICDH) (green) and OPPP enzymes (pink for cytosolic; purple for plastidial). All OPPP enzymes have identical fluxes and the flux for 6-phosphogluconate dehydrogenase (6PGDH) is shown above. Lowercase letters c and p in parenthesis stand for cytosolic and plastidial respectively. The orange vertical dashed and solid lines represented Point A and LCP respectively.

## Discussion

### The Kok method overestimates day respiration

The metabolism and regulation of day respiration is fairly complex due to its interaction with photosynthesis and photorespiration. There had been many experimental studies using various methods to study and to understand the origin of the Kok effect in C_3_ leaf previously. Till today, it remained a challenging task to determine the rate of day respiration accurately and conveniently and to understand its mechanism due to the presence of various compounding factors, such as environmental factors or cellular states. In this study, we tried to overcome these challenges by using a computational approach, constraint-based modelling, and modelled the Kok effect successfully based on a purely stoichiometry model.

Our model agrees with many observations previously reported. Not only were we able to observe the typical change in linearity in the light response curve at LCP, we also found another more subtle break in the predicted light response curve at a much lower photon flux density (Point A), resulting in the light response curve having 3 distinct phases where phases 1 (0 μmol m^−2^ s^−1^ – Point A) and 2 (Point A – LCP) occurred in stage 1 and phase 3 (LCP and above) was defined as stage 2. Using the Kok method, we determine *R*_d_ to be 1.32 μmol m^−2^ s^−1^ from the intercept of the linear fitted line above the LCP from the predicted light response curve. However, by looking at the non-photorespiratory CO_2_ evolving reactions in the flux solutions from constraint-based modelling, day respiration was found to be a constant 0.07 μmol m^−2^ s^−1^ after LCP given the same set of modelling constraints, suggesting that the Kok method could vastly overestimate day respiration (*R*_d_), which is commonly defined as the non-photorespiratory CO_2_ production during the day. Besides the magnitude of day respiration, we were able to identify the contributions of different metabolic reactions in day respiration, with major contributions from isocitrate dehydrogenase (44%) and mitochondrial pyruvate dehydrogenase (33%) (Fig. 4b).

### Potential in improving metabolic efficiency under low light

Our model overestimated the quantum yield in stage 1 (from no light to LCP) compared to the experimental measured quantum yield. Given that we applied an efficiency-based objective function in our modelling, our results suggest that plant metabolism under very low light may not be regulated in a way that is optimised for energetic efficiency. Hence our model predictions suggest that there could be room for improvement with regards to energetic efficiency in low light, though this might be due to inefficiencies in light capture and/or might come with a potential cost at the regulation level (e.g. speed and cost of regulation), both of which our model does not capture. Our model predicted an optimal set of changes in metabolic flux modes under low light assuming maximum efficiency in carbon and energy use. Unsurprisingly, our model predicted that the main metabolic processes with changes in fluxes from darkness to low light were the OPPP, the TCA cycle and lower glycolysis. An overview of the interactions and functions of these processes were summarised in Fig. 9. Some enzymes in these processes were predicted to become inactive above certain light intensities. For example, the TCA cycle was predicted to operate in a non-cyclic mode at light intensity above Point A with no flux through the reactions involving the generation of fumarate from oxoglutarate (Fig. S3), which is consistent with the experimental results obtained on *Xanthium strumarium* leaves using isotopic methods (Tcherkez et al., 2009).

**Fig. 9.**
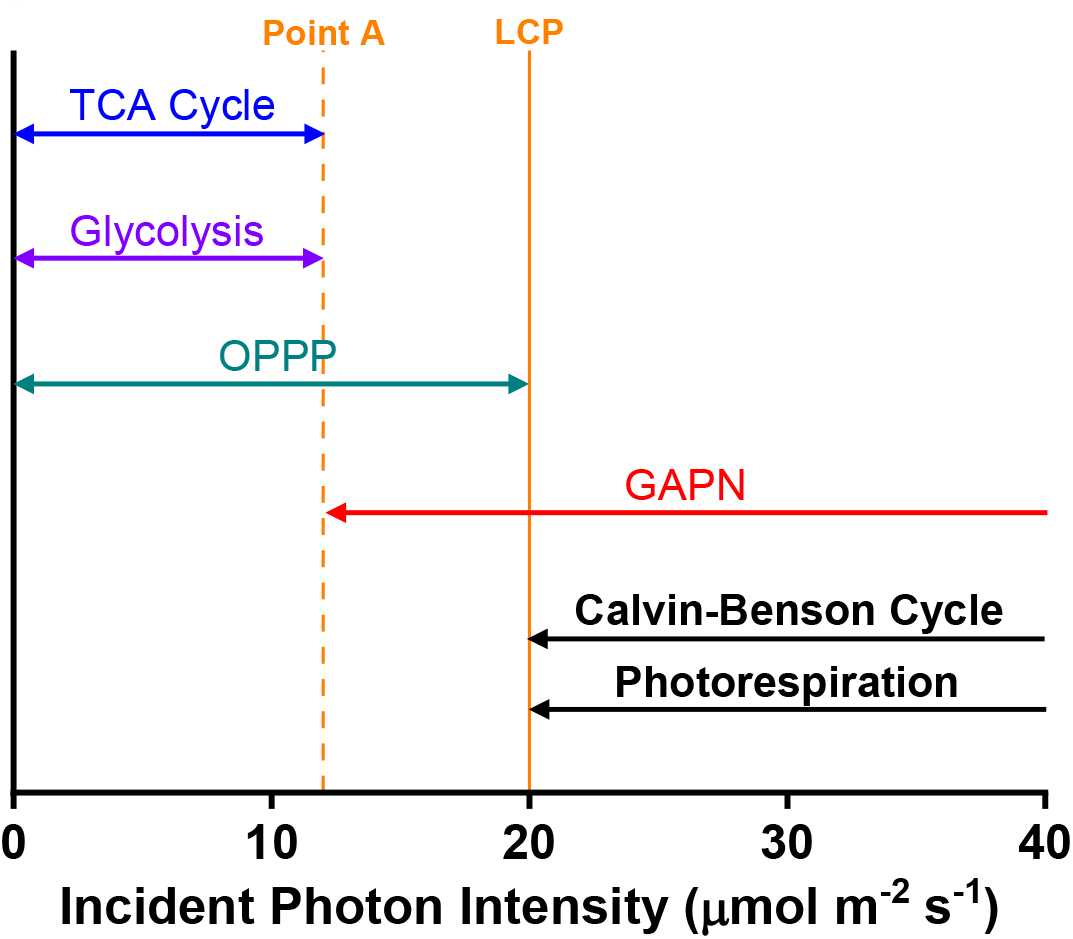
An overview of the predicted activities of metabolic processes found to be associated with the Kok effect at the various photon intensities. Enzymes in the TCA cycle and lower glycolysis pathway were predicted to be upregulated or active from darkness to Point A to produce ATP while enzymes in the OPPP were predicted to be active from darkness to the LCP to produce NADPH. Glyceraldehyde 3-phosphate dehydrogenase (NADP^+^) (GAPN) was predicted to be active at light intensities above Point A to produce NAPDH. The Calvin-Benson cycle and photorespiration were predicted to only be active at light intensities above the LCP under the energetically optimal assumption. Point A and LCP are represented by an orange vertical dashed and solid lines respectively.

From our model prediction, the light response curve can be divided into 3 distinct phases. In the phase from no light to Point A, the model predicted of the operation of the OPPP, the TCA cycle and lower glycolysis and they functioned to supply the NADPH and ATP needed in the dark. The synthesis of ATP by lower glycolysis and the TCA cycle only occurred in this phase as the fluxes of the enzymes either plateaued off or became zero after Point A. While the complete TCA cycle flux was only active in phase 1, it was interesting to see that the mitochondrial electron transport chain was predicted to be active at all times to produce the ATP needed by the leaf (Fig. S5). The mitochondrial electron transport chain flux dropped gradually in stage 1 before taking off after LCP in stage 2. This was likely to due to the combination of the decrease in flux in the TCA cycle and the increase in reductant shuttling from the plastid to the mitochondrion as light intensity increased.

From Point A to LCP, the production of NADPH was continued by the OPPP enzymes in the cytosol (Fig. 8). On top of that, GAPN was also activated in this period and began its production of NADPH needed for maintenance. Examining the predicted fluxes that were affected by NADPH demand together, i.e. OPPP enzymes, GAPN and IDH, had allowed us to understand the optimal regulation of these enzymes for energy efficiency during the transition from dark to low light (Fig. 8). As light intensity increased from 0 μmol m^−2^ s^−1^, NADPH can start to be produced in the plastid by the photosynthetic light reaction, leading to a decreased in OPPP flux in the plastid. Same applies to ATP where the TCA cycle flux decreased as ATP can start to be produced by the photosynthetic light reaction. The side effect of a decreased TCA cycle flux is a decrease in the net NADPH production by ICDH. As a result, the OPPP enzymes in the cytosolic OPPP was predicted to increase its NADPH production from darkness to Point A to compensate for the dropped in NADPH production from the TCA cycle. As light intensity increased beyond Point A, the TCA cycle was predicted to operate in a non-cyclic mode with very low flux. The production of NADPH from the OPPP is an energetically costly process as it requires the degradation of starch and produces CO_2_, hence, as light intensity increased from Point A, the model predicted a decrease in the cytosolic OPPP flux as more NADPH can be produced form the photosynthetic light reactions in the plastid. NADPH in the plastid can be shuttled to the cytosol using GAPN, resulting in an increase in GAPN flux and a decrease in OPPP flux as light intensity increased from Point A. These predictions gave us insights into the regulation of central carbon metabolism for optimal energetics at low light level.

### Mechanistic understanding of the Kok effect from constraint-based modelling

Our modelling results illustrated a number of factors that affect the light response curve, including RuBisCO carboxylation/oxygenation ratio and the magnitude of cellular maintenance costs. An increase in RuBisCO carboxylation/oxygenation ratio, i.e. reduced photorespiration, led to an increase in quantum yield in stage 2 and an apparent increase in *R*_d_ estimated by the Kok method. RuBisCO was only predicted to be active at light intensity higher than the LCP, which means the change in RuBisCO carboxylation/oxygenation ratio has no effect on the quantum yield of stage 1 and *R*_n_. Experimental studies reported that the Kok effect disappeared at low oxygen (Ishii et al., 1979; Sharp et al., 1984) or at extremely high CO_2_ concentration (Björkman and Demmig, 1987; Evans, 1987). The relative difference between the quantum yield in the dark and day had also been reported to decrease at high CO_2_ concentration (Björkman and Demmig, 1987; Evans, 1987). Our modelling results are consistent with these observations as low oxygen or high CO_2_ leads to an increase in RuBisCO carboxylation/oxygenation ratio, which in turn increases the quantum yield of stage 2 without affecting stage 1, resulting in a decrease in the relative difference between the quantum yields of stage 1 and stage 2. At extremely high CO_2_ or low oxygen level, the ratio of RuBisCO carboxylation/oxygenation activity could be high enough to cause the Kok effect to disappear as the quantum yield of stage 2 reaches that of stage 1.

*R*_d_, *R*_n_, and LCP were predicted to increase as cellular maintenance costs increased (Fig. 6). This aligns with the results reported by Ishii and Schmid Georg (2014) who reported a temperature dependence of the Kok effect in Tobacco where the *R*_d_ of Tobacco increased as temperature increased. As temperature increases, cellular maintenance costs are expected to increase due to the change in fluidity and permeability of membranes and production of reactive oxygen species (Niu and Xiang, 2018), and thus resulting in a higher day respiration. However, the quantum yields of stage 1 and stage 2 were predicted to remain largely unchanged as cellular maintenance costs varied, suggesting that the Kok effect can be observed regardless of the magnitude of maintenance costs.

Our modelling results provided a mechanistic understanding of the source of the Kok effect. The Kok effect, defined as the change in quantum yield near the LCP, could be explained by changes in flux modes involving processes in central carbon metabolism. The subtle change in quantum yield around Point A was associated with the decrease in fluxes through glycolysis and the TCA cycle from darkness to Point A. The larger change in quantum yield at the LCP was associated with the inactivation of the OPPP and the activation of the Calvin-Benson cycle at the LCP as light intensity increased. Our results support the idea that there are multiple origins of the Kok effect including a catabolic origin and a photorespiratory origin (Tcherkez et al., 2017b) as it is the change in flux mode between catabolism and photorespiration that is associated with the change in quantum yield around the LCP. Based on this reasoning, our model was able to explain the effect of CO_2_ and O_2_ mole fractions on *R*_d_ given their effect on the quantum yield of stage 2.

Being a purely stoichiometric model, our model cannot directly model several factors that had been speculated to cause the Kok effect, such as the effects of temperature (Dungan et al., 2003; Atkin et al., 2005; Ishii and Schmid Georg, 2014) or the cyclic electron flow (Laisk et al., 2005; Kou et al., 2013; Sunil et al., 2019). While we cannot rule out the contributions of these factors in the cause of the Kok effect, our model allowed us to show that the Kok effect can be explained solely from changes in flux modes between catabolism and photorespiration without any kinetic parameters.

## Materials and Methods

### Model construction

To create a suitable metabolic model to simulate the Kok effect in a C_3_ leaf in the day time, we extracted the day metabolic network from a core diel flux balance metabolic model used for modelling C_3_ and CAM leaves (Shameer et al., 2018). After extracting the day-time reactions from the core model, the reactions were then curated and modified to ensure that they are accurate and suitable for modelling the Kok effect. The complete list of modification is available in the Supplemental Table S1. The final metabolic model, which contains 645 reactions and 556 metabolites, is available in SBML and Excel formats (Supplemental Files S1 and S2).

### Model constraints and simulations

Flux balance analysis (FBA), a constraint-based mathematical modelling approach (Orth et al., 2010), was used to predict the flux distributions in the C_3_ plant metabolic model. To model the metabolism of a mature leaf, the model was constrained and fixed to export sucrose and amino-acids to the phloem at 0.78 μmol m^−2^ s^−1^ (Shameer et al., 2018) as it had been reported that the average flow velocity was independent of incident light intensity and it remained relatively constant throughout the day and night for a number of different plant species (Peuke et al., 2001; Peuke et al., 2015; Průšová, 2016). To set up the model, we had applied the constraints that was used in the C_3_ plant modelling in Shameer et al. (2018) (Supplemental Table S2). Notably, the ATP maintenance costs was fixed at 8.5 μmol m^−2^ s^−1^ as in Shameer et al. (2018). The ratio of ATP to NADPH maintenance was set to 3:1 with NADPH maintenance costs distributed equally between the cytosol, mitochondrion and chloroplast based on Cheung et al. (2013). To model the Kok effect, we ran the model simulations with varying photon intensities and allowed starch to be accumulated and degraded based on the energetic demands of the metabolic system. The primary objective function of maximising starch accumulation (minimising starch degradation) was used with the secondary objective of minimising the absolute sum of fluxes. The scripts for the model simulations can be found in Supplemental File S3. The full flux predictions of all scenarios modelled can be found in Supplemental File S4.

## Supplemental Data

Supplemental Table S1. Manual curation log of the core metabolic model.

Supplemental Table S2. Common constraints applied in all simulations.

Supplemental Figure S1. Predicted light response curves of O_2_ production of a C_3_ leaf under different incident photon intensities.

Supplemental Figure S2. Fluxes of ‘Others’ reactions involved in the production of CO_2_.

Supplemental Figure S3. Fluxes of the tricarboxylic acid (TCA) cycle enzymes in the mitochondrion.

Supplemental Figure S4. Lower glycolysis enzymatic fluxes in the cytosol at different incident photon intensities.

Supplemental Figure S5. Enzymatic fluxes of mitochondrial electron transport chain at different incident photon intensities.

Supplemental File S1 Core plant metabolic model in SBML format.

Supplemental File S2 Core plant metabolic model in Excel format.

Supplemental File S3 Scripts for model simulation of the Kok effect.

Supplemental File S4 Full flux predictions of all scenarios modelled.

## Acknowledgements

We would like to thank Yale-NUS College for supporting this research and Kristoforus B. Odang, Ayrton San Joaquin and Sewen Thy for the technical help with the modelling software.

## References

Abadie C, Boex-Fontvieille ERA, Carroll AJ, Tcherkez G (2016) *In vivo* stoichiometry of photorespiratory metabolism. Nature Plants 2:15220

Atkin OK, Bruhn D, Tjoelker MG (2005) Response of plant respiration to changes in temperature: mechanisms and consequences of variations in Q_10_ values and acclimation. *In* H Lambers, M Ribas-Carbo, eds, Plant Respiration: From Cell to Ecosystem. Springer Netherlands, Dordrecht, pp 95–135

Atkin OK, Westbeek M, Cambridge ML, Lambers H, Pons TL (1997) Leaf respiration in light and darkness (A comparison of slow- and fast-growing poa species). Plant Physiology 113:961–965

Björkman O, Demmig B (1987) Photon yield of O_2_ evolution and chlorophyll fluorescence characteristics at 77K among vascular plants of diverse origins. Planta 170:489–504

Buckley TN, Adams MA (2011) An analytical model of non-photorespiratory CO_2_ release in the light and dark in leaves of C_3_ species based on stoichiometric flux balance. Plant, Cell & Environment 34:89–112

Budde RJ, Randall DD (1990) Pea leaf mitochondrial pyruvate dehydrogenase complex is inactivated in vivo in a light-dependent manner. Proceedings of the National Academy of Sciences 87:673–676

Cheung CYM, Poolman MG, Fell DA, Ratcliffe RG, Sweetlove LJ (2014) A diel flux balance model captures interactions between light and dark metabolism during day-night cycles in C_3_ and Crassulacean Acid Metabolism leaves. Plant Physiology 165:917–929

Cheung CYM, Ratcliffe RG, Sweetlove LJ (2015) A method of accounting for enzyme costs in flux balance analysis reveals alternative pathways and metabolite stores in an illuminated Arabidopsis leaf. Plant Physiology 169:1671–1682

Cheung CYM, Williams TCR, Poolman MG, Fell DA, Ratcliffe RG, Sweetlove LJ (2013) A method for accounting for maintenance costs in flux balance analysis improves the prediction of plant cell metabolic phenotypes under stress conditions. The Plant Journal 75:1050–1061

Dungan RJ, Whitehead D, Duncan RP (2003) Seasonal and temperature dependence of photosynthesis and respiration for two co-occurring broad-leaved tree species with contrasting leaf phenology. Tree Physiology 23:561–568

Evans J (1987) The dependence of quantum yield on wavelength and growth irradiance. Functional Plant Biology 14:69–79

Farquhar GD, Busch FA (2017) Changes in the chloroplastic CO_2_ concentration explain much of the observed Kok effect: a model. New Phytologist 214:570–584

Farr TJ, Huppe HC, Turpin DH (1994) Coordination of chloroplastic metabolism in N-limited *Chlamydomonas reinhardtii* by redox modulation (I. The activation of phosphoribulosekinase and glucose-6-phosphate dehydrogenase is relative to the photosynthetic supply of electrons). Plant Physiology 105:1037–1042

Hikosaka K, Noguchi K, Terashima I (2016) Modeling leaf gas exchange. *In* K Hikosaka, Ü Niinemets, NPR Anten, eds, Canopy Photosynthesis: From Basics to Applications. Springer Netherlands, Dordrecht, pp 61-100

Hodges M, Dellero Y, Keech O, Betti M, Raghavendra AS, Sage R, Zhu X-G, Allen DK, Weber APM (2016) Perspectives for a better understanding of the metabolic integration of photorespiration within a complex plant primary metabolism network. Journal of Experimental Botany 67:3015–3026

Huppe HC, Farr TJ, Turpin DH (1994) Coordination of chloroplastic metabolism in N-limited *Chlamydomonas reinhardtii* by redox modulation (II. Redox modulation activates the oxidative pentose phosphate pathway during photosynthetic nitrate assimilation). Plant Physiology 105:1043–1048

Illeris L, Jonasson S (1999) Soil and plant CO_2_ emission in response to variations in soil moisture and temperature and to amendment with nitrogen, phosphorus, and carbon in Northern Scandinavia. Arctic, Antarctic, and Alpine Research 31:264–271

Ishii R, Schmid Georg H (2014) The Kok effect and its relationship to photorespiration in Tobacco. In Zeitschrift für Naturforschung C, Vol 36, p 450

Ishii R, Shibayama M, Murata Y (1979) Effect of light on the CO_2_ evolution of C_3_ and C_4_ plant in relation to the Kok effect. Japanese Journal of Crop Science 48:52–57

Ishii R, Takehara T, Murata Y, Miyachi S (1977) Effects of light intensity on the rates of photosynthesis and photorespiration in C_3_ and C_4_ plants. In A Mitsui, S Miyachi, A San Pietro, S Tamura, eds, Biological Solar Energy Conversion. Academic Press, pp 265–271

Kok B (1948) A critical consideration of the quantum yield of *Chlorella* photosynthesis. Enzymologia 13: 1–56

Kok B (1949) On the interrelation of respiration and photosynthesis in green plants. Biochimica et Biophysica Acta 3:625–631

Kok B (1956) On the inhibition of photosynthesis by intense light. Biochimica et Biophysica Acta 21:234–244

Kou J, Takahashi S, Oguchi R, Badger MR, Chow WS (2013) Quantification of cyclic electron flow in spinach leaf discs. In. Springer Berlin Heidelberg, Berlin, Heidelberg, pp 271–274

Kwon H, Park TY, Hong J, Lim JH, Kim J (2009) Seasonality of net ecosystem carbon exchange in two major plant functional types in Korea. Asia-Pacific Journal of Atmospheric Sciences 45:149–163

Laisk A, Eichelmann H, Oja V, Peterson RB (2005) Control of cytochrome b_6_f at low and high light intensity and cyclic electron transport in leaves. Biochimica et Biophysica Acta (BBA) - Bioenergetics 1708:79–90

Lee CP, Eubel H, Millar AH (2010) Diurnal changes in mitochondrial function reveal daily optimization of light and dark respiratory metabolism in *Arabidopsis*. Molecular & Cellular Proteomics 9:2125–2139

Lin M, Turpin DH, Plaxton WC (1989) Pyruvate kinase isozymes from the green alga, *Selenastrum minutum*: II. Kinetic and regulatory properties. Archives of Biochemistry and Biophysics 269:228–238

Niu Y, Xiang Y (2018) An overview of biomembrane functions in plant responses to high-temperature stress. Frontiers in Plant Science 9:915–915

Nordstroem C, Soegaard H, Christensen TR, Friborg T, Hansen BU (2001) Seasonal carbon dioxide balance and respiration of a high-arctic fen ecosystem in NE-Greenland. Theoretical and Applied Climatology 70:149–166

Orth JD, Thiele I, Palsson BØ (2010) What is flux balance analysis? Nature Biotechnology 28:245–248

Peuke AD, Gessler A, Trumbore S, Windt CW, Homan N, Gerkema E, Van as H (2015) Phloem flow and sugar transport in *Ricinus communis* L. is inhibited under anoxic conditions of shoot or roots. Plant, Cell & Environment 38:433–447

Peuke AD, Rokitta M, Zimmermann U, Schreiber L, Haase A (2001) Simultaneous measurement of water flow velocity and solute transport in xylem and phloem of adult plants of *Ricinus communis* over a daily time course by nuclear magnetic resonance spectrometry. Plant, Cell & Environment 24:491–503

Průšová A (2016) Light on phloem transport (an MRI approach). PhD thesis. Wageningen University & Research, Wageningen, NL

Scheible W-R, Krapp A, Stitt M (2000) Reciprocal diurnal changes of phosphoenolpyruvate carboxylase expression and cytosolic pyruvate kinase, citrate synthase and NADP-isocitrate dehydrogenase expression regulate organic acid metabolism during nitrate assimilation in tobacco leaves. Plant, Cell & Environment 23:1155–1167

Shameer S, Baghalian K, Cheung CYM, Ratcliffe RG, Sweetlove LJ (2018) Computational analysis of the productivity potential of CAM. Nature Plants 4:165–171

Sharp RE, Matthews MA, Boyer JS (1984) Kok effect and the quantum yield of photosynthesis: light partially inhibits dark respiration. Plant Physiology 75:95–101

Singh KK, Chen C, Gibbs M (1993) Photoregulation of fructose and glucose respiration in the intact chloroplasts of *Chlamydomonas reinhardtii* F-60 and spinach. Plant Physiology 101:1289–1294

Sunil B, Saini D, Bapatla RB, Aswani V, Raghavendra AS (2019) Photorespiration is complemented by cyclic electron flow and the alternative oxidase pathway to optimize photosynthesis and protect against abiotic stress. Photosynthesis Research 139:67–79

Tcherkez G, Boex-Fontvieille E, Mahé A, Hodges M (2012) Respiratory carbon fluxes in leaves. Current Opinion in Plant Biology 15:308–314

Tcherkez G, Gauthier P, Buckley TN, Busch FA, Barbour MM, Bruhn D, Heskel MA, Gong XY, Crous K, Griffin KL, Way DA, Turnbull MH, Adams MA, Atkin OK, Bender M, Farquhar GD, Cornic G (2017a) Tracking the origins of the Kok effect, 70 years after its discovery. New Phytologist 214:506–510

Tcherkez G, Gauthier P, Buckley TN, Busch FA, Barbour MM, Bruhn D, Heskel MA, Gong XY, Crous KY, Griffin K, Way D, Turnbull M, Adams MA, Atkin OK, Farquhar GD, Cornic G (2017b) Leaf day respiration: low CO_2_ flux but high significance for metabolism and carbon balance. New Phytologist 216:986–1001

Tcherkez G, Mahé A, Gauthier P, Mauve C, Gout E, Bligny R, Cornic G, Hodges M (2009) In folio respiratory fluxomics revealed by ^13^C isotopic labeling and H/D isotope effects highlight the noncyclic nature of the tricarboxylic acid “cycle” in illuminated leaves. Plant Physiology 151:620–630

Tovar-Méndez A, Miernyk JA, Randall DD (2003) Regulation of pyruvate dehydrogenase complex activity in plant cells. European Journal of Biochemistry 270:1043–1049

Turnbull MH, Whitehead D, Tissue DT, Schuster WSF, Brown KJ, Griffin KL (2001) Responses of leaf respiration to temperature and leaf characteristics in three deciduous tree species vary with site water availability. Tree Physiology 21:571–578

Wilson KB, Baldocchi DD, Hanson PJ (2001) Leaf age affects the seasonal pattern of photosynthetic capacity and net ecosystem exchange of carbon in a deciduous forest. Plant, Cell & Environment 24:571–583

